# Replicability, repeatability, and long-term reproducibility of cerebellar morphometry

**DOI:** 10.1101/2020.09.02.279786

**Authors:** Peter Sörös, Louise Wölk, Carsten Bantel, Anja Bräuer, Frank Klawonn, Karsten Witt

## Abstract

To identify robust and reproducible methods of cerebellar morphometry that can be used in future large-scale structural MRI studies, we investigated the replicability, repeatability, and longterm reproducibility of three fully-automated software tools: FreeSurfer, CERES, and ACAPULCO. Replicability was defined as computational replicability, determined by comparing two analyses of the same high-resolution MRI data set performed with identical analysis software and computer hardware. Repeatability was determined by comparing the analyses of two MRI scans of the same participant taken during two independent MRI sessions on the same day for the Kirby-21 study. Long-term reproducibility was assessed by analyzing two MRI scans of the same participant in the longitudinal OASIS-2 study. We determined percent difference, the image intraclass correlation coefficient, the coefficient of variation, and the intraclass correlation coefficient between two analyses. Our results show that CERES and ACAPULCO use stochastic algorithms that result in surprisingly high differences between identical analyses for ACAPULCO and small differences for CERES. Changes between two consecutive scans from the Kirby-21 study were less than ±5% in most cases for FreeSurfer and CERES (i.e., demonstrating high repeatability). As expected, long-term reproducibility was lower than repeatability for all software tools. In summary, CERES is an accurate, as demonstrated before, and reproducible tool for fully-automated segmentation and parcellation of the cerebellum. We conclude with recommendations for the assessment of replicability, repeatability, and longterm reproducibility in future studies on cerebellar structure.

## Introduction

Physiology and pathophysiology of the cerebellum have received growing attention in basic and clinical neurosciences (1–3). Early nineteenth century neuroscientists, especially Luigi Rolando and Pierre Flourens, have established the crucial role of the cerebellum in motor control (4) and, more specifically, motor coordination (5). More recently, the role of motor learning (6, 7) and the non-motor functions of the cerebellum (8) have been investigated in greater detail. The cerebellar contributions to various cognitive (9) and emotional functions (10) as well as timing (11, 12) have been acknowledged. Moreover, structural changes of the cerebellum in healthy aging (13) and neurodegenerative disease (14, 15) have been studied.

The advent of magnetic resonance imaging (MRI) has opened the door to quantitative, non-invasive investigations of cerebellar morphology. The segmentation of the cerebellum into gray and white matter and the parcellation into lobes and single lobules turned out to be challenging because of its tightly folded structure, consisting of numerous small folia, the equivalent of cerebral gyri. Moreover, the anatomy of the cerebellum is characterized by pronounced inter-individual differences (16, 17). Manual slice-by-slice labeling of MRIs by an expert neuroanatomist is considered the gold standard of cerebellar research (18). Nevertheless, manual segmentation and parcellation have major disadvantages, requiring expert knowledge and being observer-dependent and time-consuming, and are not feasible in large-scale studies.

To overcome the limitations of manual identification of cerebellar structures, several fully automated methods for cerebellar morphometry have been developed and made publicly available (for a review, see Carass et al. (19)). The results of several of these methods have been compared with manually labeled adult and pediatric cerebellar data sets (19). In this comparison, an improved version of the patch-based multiatlas segmentation tool CERES (CEREbellum Segmentation) (20) exhibited highest accuracy and outperformed established methods, such as the MATLAB toolbox SUIT (Spatially Unbiased Infra-tentorial Template) (16, 21). While the accuracy of CERES and other methods have been established, the reproducibility of fully automated cerebellar morphometry has not been determined so far.

In the present study we investigate the replicability, repeatability, and long-term reproducibility of cerebellar morphometry using three independent MRI data sets and three software packages based on different computational approaches. The definitions of replicability, repeatability, and reproducibility follow the suggestions by Nichols et al. (22). Replicability is defined as computational or analysis replicability, determined by comparing two analyses of the same MRI data set performed with identical analysis software and computer hardware. Repeatability is determined by comparing the analyses of two MRI scans of the same participant taken during two independent MRI sessions on the same day. Long-term reproducibility, finally, is assessed by analyzing two MRI scans of the same participant in a longitudinal study. We decided to test the following three software packages: (1) FreeSurfer, an established and widely used approach of subcortical segmentation, based on a probabilistic atlas, which performs cerebellar segmentation, but not parcellation (23), (2) CERES, a recent segmentation and parcellation method based on a multiatlas label fusion technique (20), the most accurate software tool in the comparison by Carass et al. (19), and (3) ACAPULCO, a very recent and promising parcellation approach based on convolutional neural networks (24), not included in the comparison by Carass et al. (19). In a separate paper, the developers of ACAPULCO demonstrated comparable accuracy of their software relative to CERES for adult data and even superior accuracy in several regions for pediatric data (24).

The ultimate aim of this study is to identify robust and reproducible methods of fully automated cerebellar morphometry that can be used in MRI studies with large sample sizes.

## Methods

### MRI data

For this study, three independent data sets of T1weighted MRIs of the entire brain have been analyzed with three different fully automated software packages: FreeSurfer 7.1.0 (23), CERES (20), and ACAPULCO (24).

#### Replicability: ChroPain2 study

To investigate the analysis replicability of cerebellar morphometry, we performed two separate, but identical analyses of high-resolution structural MRIs of 23 healthy individuals (17 women, 6 men) who served as control participants for the ChroPain2 study. Inclusion and exclusion criteria have been published previously (25). Mean age ± standard deviation was 51 ± 10 years (minimum: 30 years, maximum: 66 years). All participants provided written informed consent for participation in this study. The study was approved by the Medical Research Ethics Board, University of Oldenburg, Germany (2017-059) and was preregistered with the German Clinical Trials Register (DRKS00012791)^1^.

MR images of the entire brain were acquired in the Neuroimaging Unit, School of Medicine and Health Sciences, University of Oldenburg^2^, on a research-only Siemens MAGNETOM Prisma whole-body scanner (Siemens, Erlangen, Germany) at 3 Tesla with a 64-channel head/neck receive-array coil. A 3-dimensional high-resolution and high-contrast T1-weighted magnetization prepared rapid gradient echo (MPRAGE) sequence was used (26). Imaging parameters were: TR (repetition time; between two successive inversion pulses): 2000 ms, TE (echo time): 2.07 ms, TI (inversion time): 952 ms, flip angle: 9°, isotropic voxel size: 0.75 × 0.75 × 0.75 mm^3^, 224 sagittal slices, k-space interpolation-based in-plane acceleration (GRAPPA) with an acceleration factor of 2 (27), time of acquisition: 6:16 min. Siemens’ prescan normalization filter was used for online compensation of regional signal inhomogeneities.

#### Repeatability: Kirby-21 study

To investigate repeatability of cerebellar morphometry, we analyzed data from the Kirby21 multi-modal MRI reproducibility study (28), performed at the F.M. Kirby Research Center for Functional Brain Imaging, Kennedy Krieger Institute, Baltimore, MD, USA. For this study, each participant received two identical MRI examinations on the same day, each consisting of several sequences, including a T1-weighted MPRAGE sequence. After the first examination, participants left the scanner room for a short break and were then repositioned and scanned with the identical imaging protocol a second time. The time interval between the two T1-weighted images was approximately 1 hour. MRIs were acquired from 21 individuals (10 women, 11 men) with no history of neurological disorders. Mean age ± standard deviation was 32 ± 9 years (minimum: 22 years, maximum: 61 years). For a detailed description of the entire study, see (28). The data set is publicly available for download^3^ and has been used in several studies on the reproducibility of MRI analyses (e.g., (29, 30)).

MR images of the entire brain were acquired at 3 Tesla using a Philips Achieva MR scanner (Philips Healthcare, Best, The Netherlands) with an 8-channel receive-array head coil. Imaging parameters for the MPRAGE sequence were: TR (between two successive gradient echoes): 6.7 ms, TE: 3.1 ms, TI: 842 ms, flip angle: 8°, voxel size: 1 × 1 × 1.2 mm^3^, image domain-based in-plane acceleration (SENSE) with an acceleration factor of 2, duration: 5:56 min.

#### Long-term reproducibility: OASIS-2 study

To investigate long-term reproducibility of cerebellar morphometry, we performed analyses of MR images acquired for the Open Access Series of Imaging Studies (OASIS-2) (31), performed at the Washington University School of Medicine, St. Louis, MO, USA. The OASIS-2 study comprises longitudinal MR examinations of patients with Alzheimer’s disease and healthy controls. For the present study of cerebellar morphometry, we analyzed the data of 72 individuals (50 women, 22 men) who remained cognitively unimpaired throughout the study, as demonstrated by a Clinical Dementia Rating (CDR) score of 0 (32). Mean age at inclusion ± SD was 75 ± 8 years (minimum: 60 years, maximum: 93 years). For the OASIS-2 study, participants received 2-5 MRI examinations months or years apart; each MRI examination consisted of 3-4 T1-weighted MRI scans. For the present study, we only considered the first two MRI examinations of each participant. If more than one MRI scan was available for one examination, we chose the first one. The mean interval ± SD between the two MRIs was 738 ± 249 days (minimum: 182, maximum: 1510 days). All MRIs were obtained with the same scanner with identical pulse sequences. For a detailed description of the study and the CDR scale, see Marcus et al. (31). OASIS-2 data sets are publicly available for download.^4^

MR images of the entire brain were acquired on a Siemens Vision whole-body scanner (Siemens, Erlangen, Germany) at 1.5 Tesla. Imaging parameters for the MPRAGE sequence were: TR (between two successive gradient echoes): 9.7 ms, TE: 4 ms, TI 20 ms, flip angle: 10°, voxel size: 1 × 1 × 1.25 mm^3^, 128 sagittal slices.

### Data analysis

FreeSurfer and ACAPULCO analyses were performed on the high-performance computer cluster CARL^5^ at the University of Oldenburg, Germany, running Red Hat Enterprise Linux. CERES was run through the online MRI Brain Volumetry System volBrain (33). CERES can only be used through the volBrain website and was not available for installation on our computer cluster. All analyses were done fully automated. Manual editing of output images was not performed, because the aim of this study was to assess reproducibility of cerebellar morphometry for future use in largescale data sets.

#### FreeSurfer

For automated analysis of subcortical structures, including the cerebellum, the FreeSurfer 7.1.0 image analysis suite was used, which is freely available for download online^6^ (36). Processing was done with the `recon-all-all` command. For the ChroPain2 and the Kirby-21 data sets, the `-3T` and `-mprage` flags were used. For the OASIS-2 data sets, the `-mprage` flag was used. Processing started with automated transformation to Talairach space, followed by intensity normalization of the output images and removal of non-brain tissue using a hybrid approach that combines watershed algorithms and deformable surface models (37). During segmentation, a neuroanatomical label is assigned to all voxels of the T1-weighted MRI based on a probabilistic atlas, derived from a manually labeled training set (23), using a Bayesian approach. Details of atlas construction, registration of the probabilistic atlas to the individual MRI, and segmentation based on the assumption that spatial distribution of labels can be approximated by an anisotropic nonstationary Markov random field are given by Fischl et al. (23). FreeSurfer reports the volumes of the left and right cerebellar cortex and the left and right cerebellar white matter (Figure 1).

**Fig. 1.**
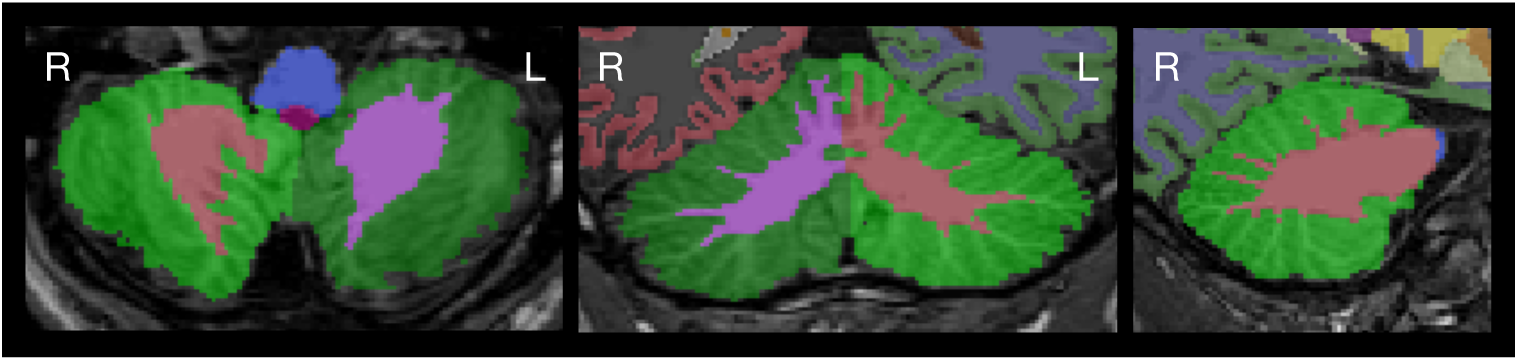
Cerebellar segmentation as determined by FreeSurfer. Images were created with FSLeyes. The left image shows a horizontal, the middle image a coronal, and the right image a sagittal section of the cerebellum. Images are in radiological convention (the left side of the cerebellum is on the right side of the image). Cerebellar cortex is displayed in green color.

Parallelization was not used, all processes were run on a single computer core of a high-performance computer cluster. Processing of the first MRI of the first MR examination of participant OAS2_0095 failed due to an error during topology correction (with and without the `-mprage` flag). We analyzed the second MRI of the first examination instead; processing finished without error.

#### CERES (CEREbellum Segmentation)

CERES is an automated pipeline for cerebellar segmentation and parcellation (20) and is part of the volBrain Automated MRI Brain Volumetry System (33). In brief, CERES receives an anonymized T1-weighted MRI brain volume in NIfTI format through the volBrain website^7^, performs image preprocessing, and labels cerebellar voxels based on Optimized Patch-Match Label fusion (38).

Preprocessing includes (1) denoising (39), (2) bias field correction using the N4 algorithm (40), (3) linear registration to the MNI152 standard space template using Advanced Normalization Tools (ANTs) (41, 42), (4) cropping of the cerebellum area, (5) non-linear registration to the cropped MNI152 template using ANTs (41, 42), and (6) local intensity normalization. Labelling of cerebellar voxels was performed with non-local patch-based label fusion, a multiatlas segmentation technique combining segmentations from multiple reference atlases, initially developed for hippocampal segmentation (43, 44). The atlases were created based on manually segmented high-resolution MR images from 5 healthy volunteers (3 women, 2 men, aged 29–57 years) (18), available for download^8^. CERES determines the entire volume, cortical thickness, and gray matter volume of all regions listed in Table 1, separately for the left and right side of the cerebellum (Figure 2). Of note, we have used the publicly available first version of CERES. All analysis steps have been determined by the developers; changes of analysis methods or parameters are not possible. In the study on accuracy of cerebellar morphometry performed by Carass et al. (19), an improved version (CERES2) was tested, which employs an improved intensity normalization method and a systematic error correction step; CERES2 has not been released for public use so far.

**Table 1.**
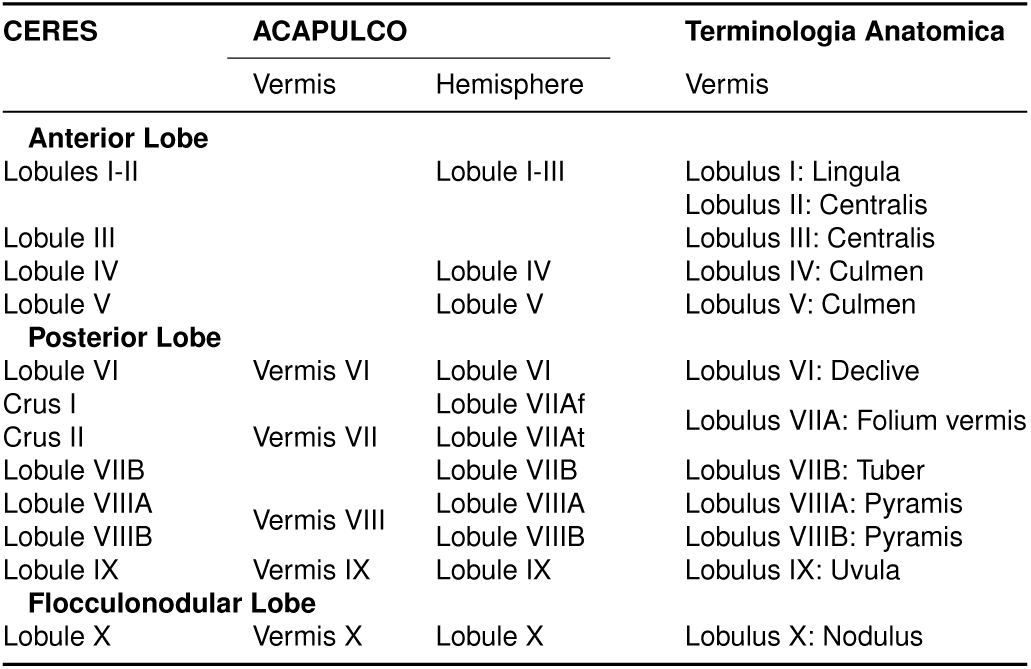
Cerebellar regions parcellated in CERES and ACAPULCO. CERES determines the entire volume (cm^3^), the mean cortical thickness (mm), and the gray matter volume (cm^3^) of each region. ACAPULCO determines the volume (mm^3^) of each region. CERES and ACAPULCO make use of the cerebellar nomenclature proposed by Schmahmann et al. (34). In addition, the traditional names of vermical regions according to the Terminologia Anatomica (35) are listed. The less common names of hemispheric regions were omitted.

**Fig. 2.**
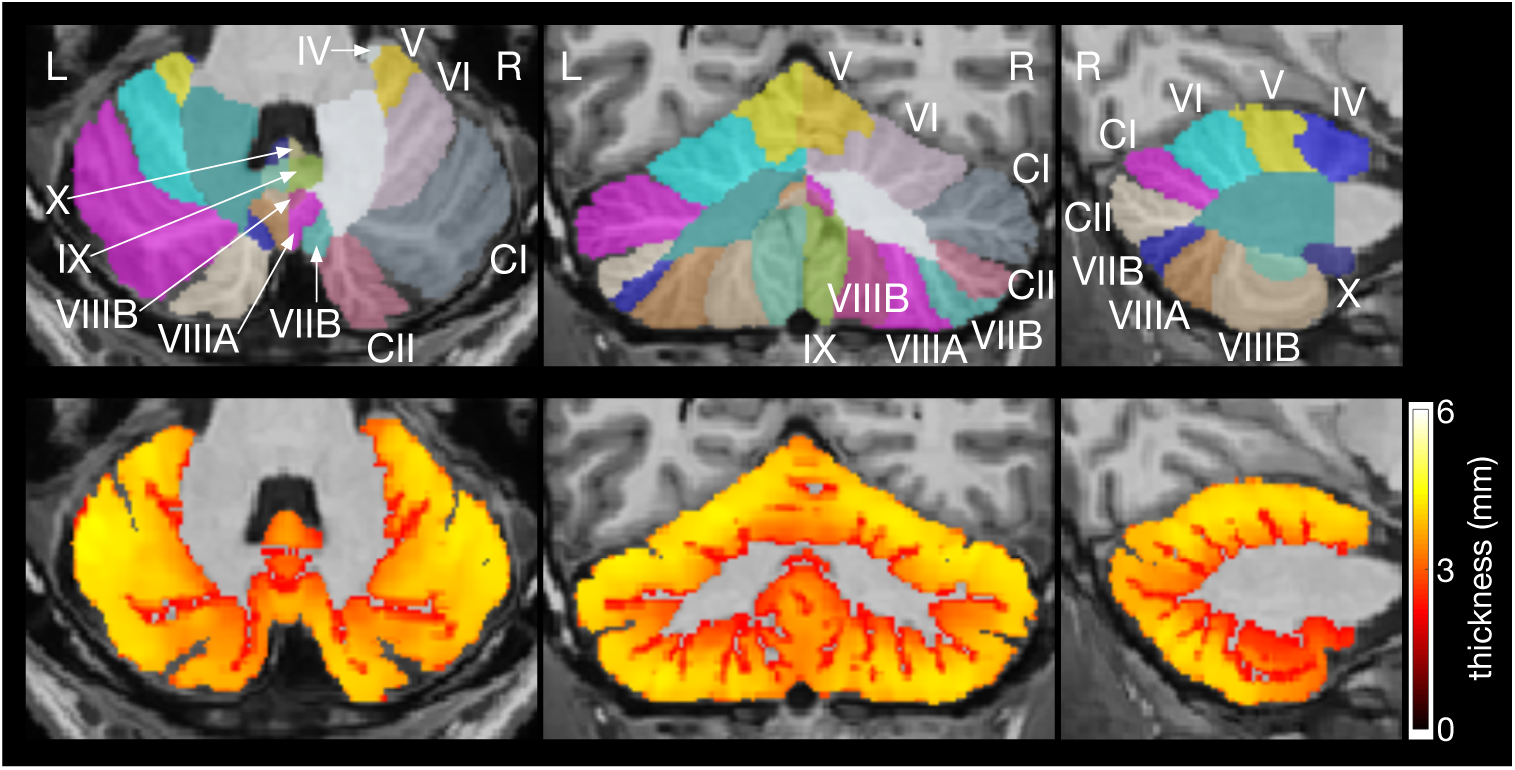
Cerebellar parcellation (upper row) and thickness of cerebellar cortex (lower row) as determined by CERES. The left images show horizontal, the middle images coronal, and the right images sagittal sections of the cerebellum. Images were produced by CERES and are in neurological convention (the left side of the cerebellum is on the left side of the image). The roman numerals of the cerebellar lobules were added. CI denotes Crus I; CII, Crus II.

#### ACAPULCO (automatic cerebellum anatomical parcellation using U-Net with locally constrained optimization)

Han et al. (24) developed a method using convolutional neural networks for cerebellar parcellation (ACAPULCO). ACA-PULCO processes T1-weighted images of the brain in NIfTI format, preferentially acquired with an MPRAGE sequence. A Singularity image of this software is publicly available^9^. We ran this image on University of Oldenburg’s HPC cluster using Singularity 2.6.

As suggested by the developers, all images were first cropped with the `robustfov` command provided by FSL^10^ to remove the lower head and neck in MRIs with large field-of-view. Processing within ACAPULCO included (1) estimation of a brain mask using Robust Brain Extraction (ROBEX)^11^ (45) for subsequent bias field correction, (2) bias field correction using the N4 algorithm (40), (3) linear registration to MNI space using the 1 mm isotropic ICBM 2009c nonlinear symmetric template^12^ using ANTs (41, 42), (4) parcellation of the cerebellum as described by Han et al. (24), and (5) transformation of the parcellation into original space using ANTs with the `MultiLabel` interpolation. For cerebellar parcellation, ACAPULCO employs two three-dimensional convolutional neural networks. First, a locating network is used to predict a bounding box around the cerebellum. Second, a parcellating network is used to parcellate the cerebellum using the entire region within the bounding box (24). ACAPULCO employs the TensorFlow software library for Python^13^ and the GNU Parallel tool (46). The cerebellar regions identified by ACAPULCO are summarized in Table 1; ACAPULCO reports the entire volume of the left and right lobules and the vermis regions in the midline (Figure 3). Of note, all analysis steps have been determined by the developers; changes of analysis methods or parameters are not possible.

**Fig. 3.**
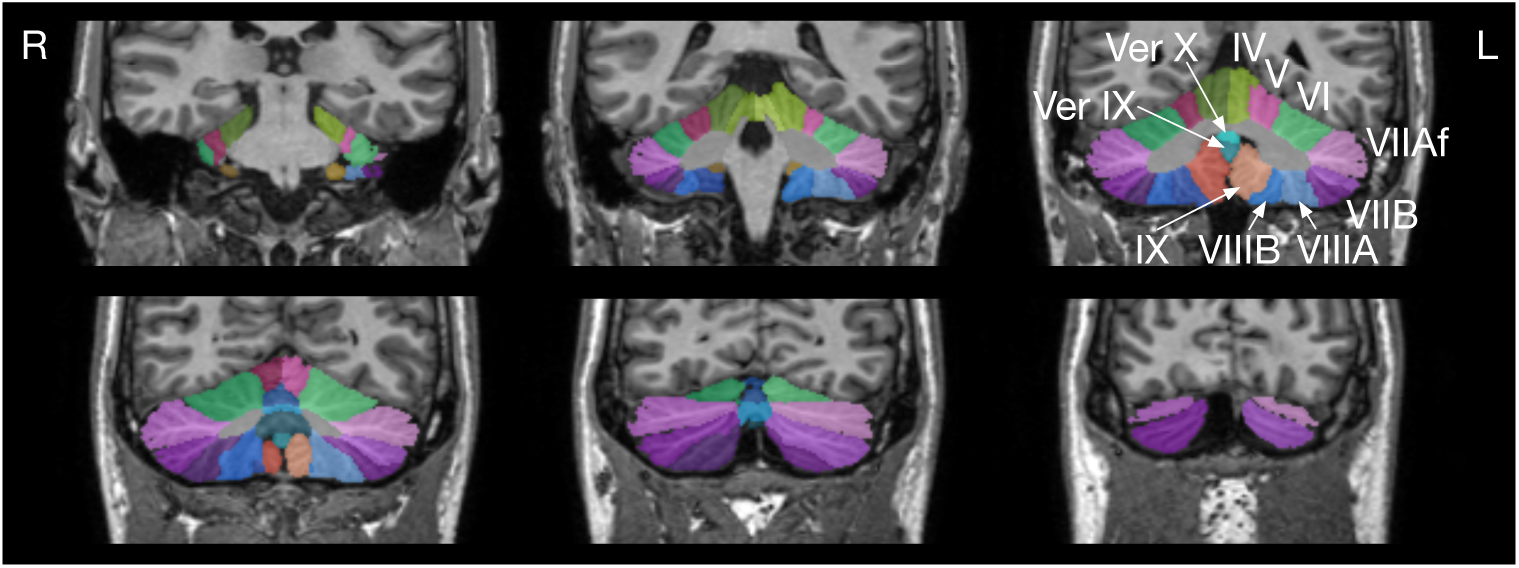
Cerebellar parcellation as determined by ACAPULCO. The upper left image is the most anterior coronal section. Images were produced by ACAPULCO. The upper part of each image was cropped and the roman numerals of the cerebellar lobules were added. Ver IX denotes Vermis IX I; Ver X, Vermis X. Images are in radiological convention (left side of the cerebellum is on the right side of the image).

Unexpectedly, we found large differences between the first and second analysis of the high-resolution ChroPain2 data set using ACAPULCO. To investigate ACAPULCO’s analysis replicability with a different data set using larger voxel sizes, we performed another two separate analyses of the T1-weighted MRIs available in the Kirby-21 study using Singularity 3.4 (in the meantime, Singularity 2.6 had been deleted from the cluster). For scan KKI2009-33, the locating network of ACAPULCO predicted an incorrect bounding box in one of these analyses, placing it well above the cerebellum, leading to erroneous results of the parcellating network. This scan was excluded from the assessment of analysis replicability in the Kirby-21 study (Figure 4C).

**Fig. 4.**
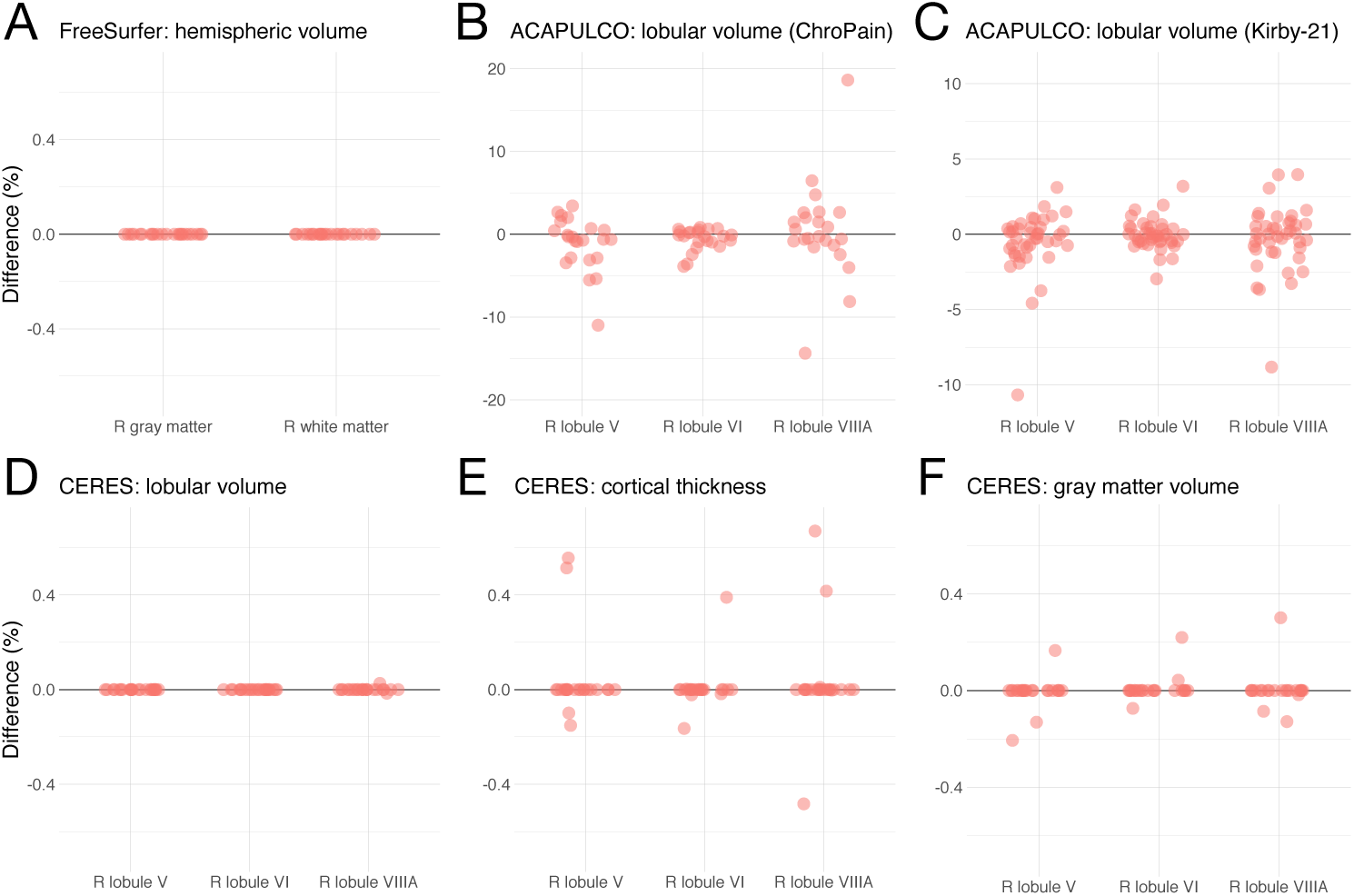
Analysis replicability of cerebellar morphometry in the ChroPain2 study using FreeSurfer (A), ACAPULCO (B), and CERES (D, E, F). Analysis replicability was also assessed with all T1-weighted MRIs of the Kirby-21 study using ACAPULCO (C). The graphs show percent difference between the first and second analysis of the same data set for right gray and white matter (FreeSurfer) and the right lobules V, VI, and VIIIA (ACAPULCO, CERES). **Note:** the scales of the y-axes differ across graphs.

### Statistical analysis

For further data analyses, we calculated the percent difference between the first and second analysis of one MRI (ChroPain2 study) or the first and second MRI (Kirby-21 and OASIS-2 studies), and determined the coefficient of variation (CV) and the intraclass correlation coefficient (ICC) for each cerebellar region. We also computed the image intraclass correlation coefficient (I2C2) (47) for cerebellar parcellations obtained by CERES.

#### Coefficient of variation

The coefficient of variation (CV) describes the level of variability within a sample independently of the absolute values of the observations. To calculate the CV, the standard deviation across all measurements of one parameter (including the results of the first and second analysis of one MRI or the analyses of the first and second MRI) was divided by the (absolute) mean across all measurements and expressed as %. In addition, the lower and upper 95% confidence intervals were estimated using R for Windows (48).

#### Intraclass correlation coefficient

The intraclass correlation coefficient (ICC) is a measure of within-subject variability relative to between-subject variability. ICC estimates and their lower and upper 95% confidence intervals were calculated using the R package *psych*^14^ and the function ICC (49). Following the suggestions of Liljequist et al. (50) we first calculated all three single-measurement ICCs (51, 52). The results of all three formulas were very similar, indicating the absence of bias (systematic error). Hence, we report the oneway random effects, absolute agreement, single measurement ICC according to McGraw and Wong (52) or the ICC(1,1) according to Shrout and Fleiss (51). ICC confidence intervals indicate poor reliability (<0.5), moderate reliability (0.5 – 0.75), good reliability (0.75 – 0.9), or excellent reliability (>0.9) (53).

#### Image intraclass correlation coefficient

The I2C2 has been developed as a global measure of reliability for imaging data (47). The (I2C2)^15^ was calculated for all cerebellar parcellations obtained by CERES for the Kirby-21 and OASIS-2 data sets using the *I2C2* package version 0.2.4 (47) for Neuroconductor (54).

First, all parcellated images created by CERES were split into 24 image files containing one parcellation only (labels 1-12 for the left cerebellum, labels 101-112 for the right cerebellum). Then, .nii files were imported into R using the `readnii` function of the *neurobase* package for Neuroconductor. Finally, the I2C2 and the nonparametrically boot-strapped 95% confidence interval of the I2C2 (with 1000 repetitions) between the first and second image of each participant were estimated.

## Results

In this section, we will visualize results of cerebellar morphometry for the right lobules V, VI, VIIIA obtained by CERES and ACAPULCO (Figures 4, 5, and 6). These lobules were chosen because of their critical role in motor and non-motor functions of the cerebellum. According to an activation likelihood estimate meta-analysis of neuroimaging studies (55), (1) right lobule V is associated with motor and somatosensory processing, (2) right lobule VI is associated with motor, spatial, language, working memory, and emotional processing, and (3) right lobule VIIIA is associated with motor and working memory processing.

**Fig. 5.**
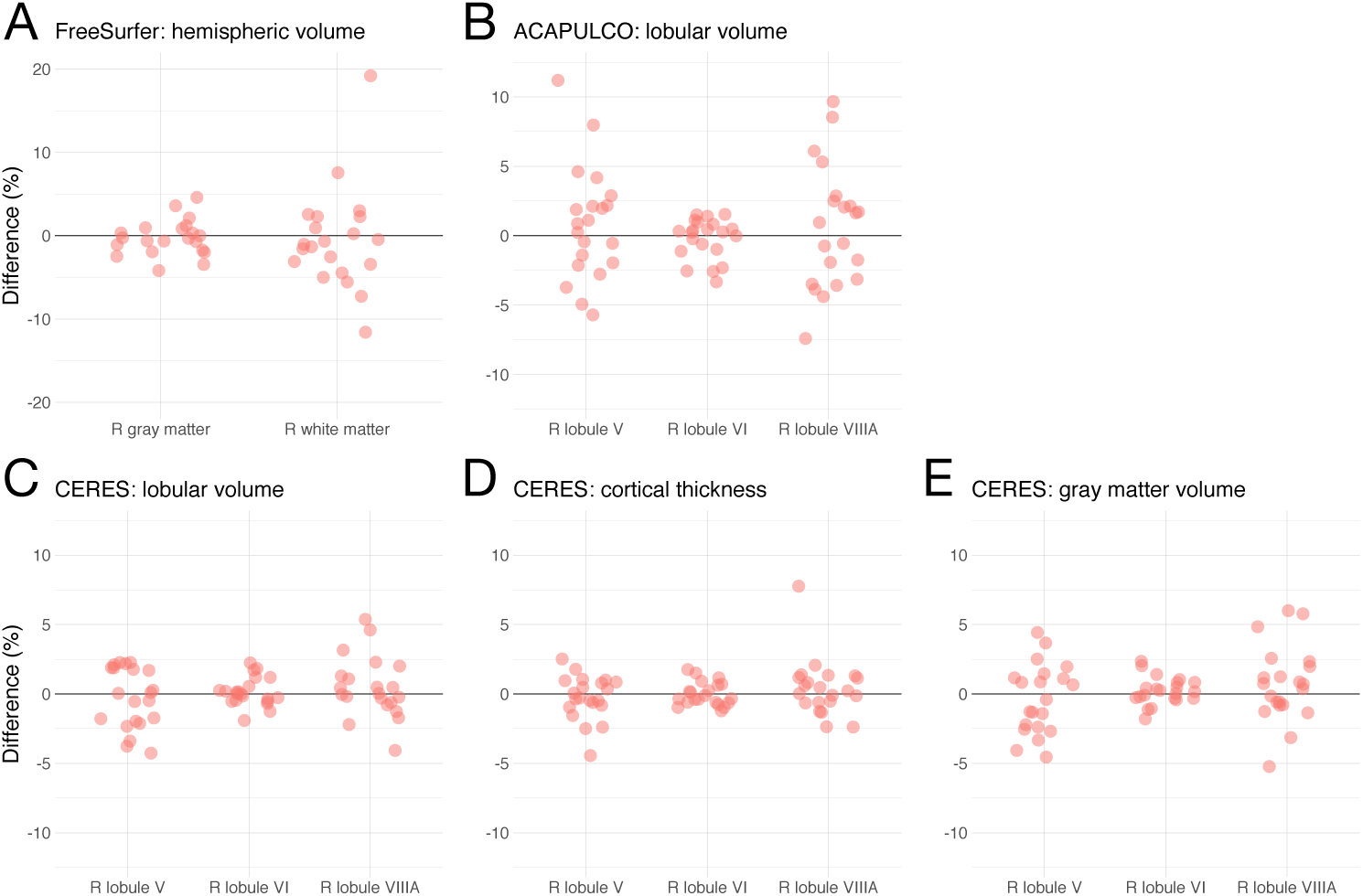
Repeatability of cerebellar morphometry in the Kirby-21 study. The graphs show percent difference between the first and second MRI acquired on the same day for right gray and white matter using FreeSurfer (A) and the right lobules V, VI, and VIIIA using ACAPULCO (B) and CERES (C, D, E). **Note:** the scales of the y-axes differ across graphs.

**Fig. 6.**
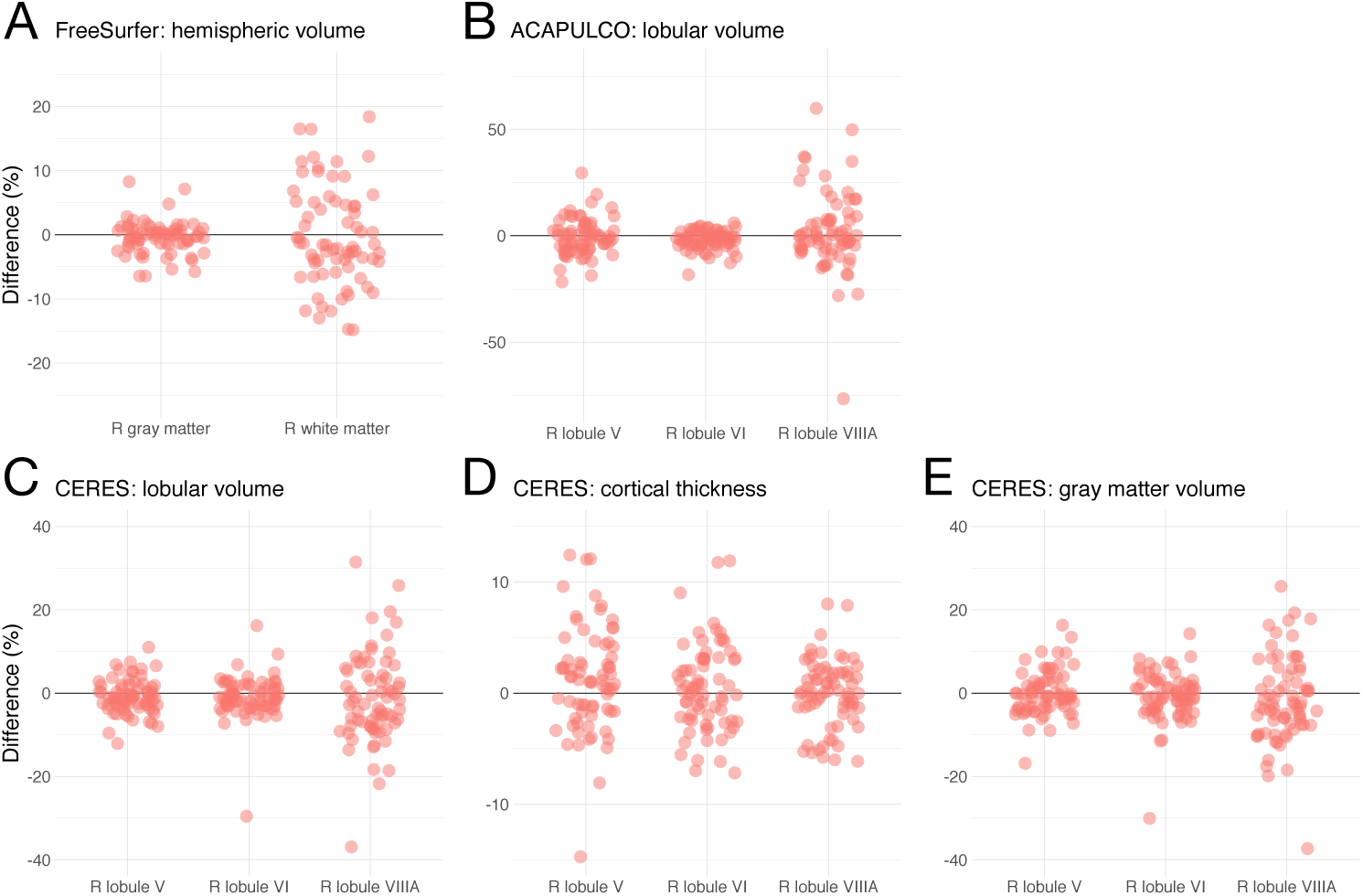
Long-term reproducibility of cerebellar morphometry in the OASIS-2 study. The graphs show percent difference between the first and second MRI for right gray and white matter using FreeSurfer (A) and the right lobules V, VI, and VIIIA using ACAPULCO (B) and CERES (C, D, E). The mean interval between the two MRIs was 738 ± 249 days (minimum: 182, maximum: 1510 days). **Note:** the scales of the y-axes differ across graphs.

We will also present the coefficient of variation (CV) and the intraclass correlation coefficient (ICC) for the results of all analyses with FreeSurfer, CERES, and ACAPULCO (Tables 3-4). For CERES parcellations, we will also provide the image intraclass correlation coefficients (I2C2) (Table 2). Supplementary data are available at the Open Science Framework (OSF)^16^.

**Table 2.**
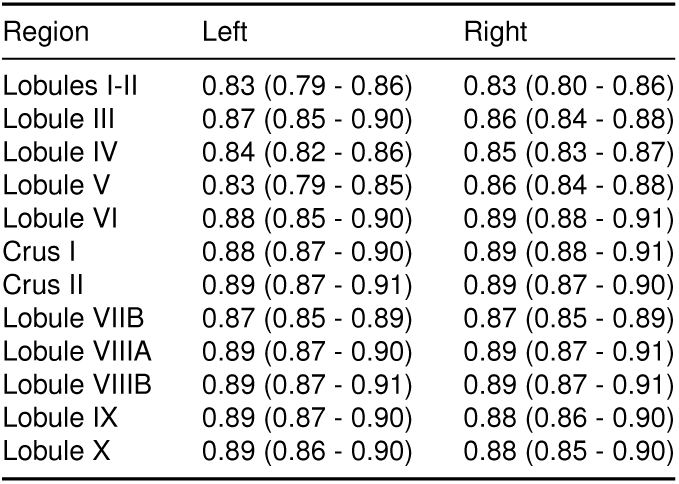
Image intraclass correlation coefficients (I2C2) and 95% confidence intervals for cerebellar regions obtained by CERES with data from the Kirby-21 study.

**Table 3.**
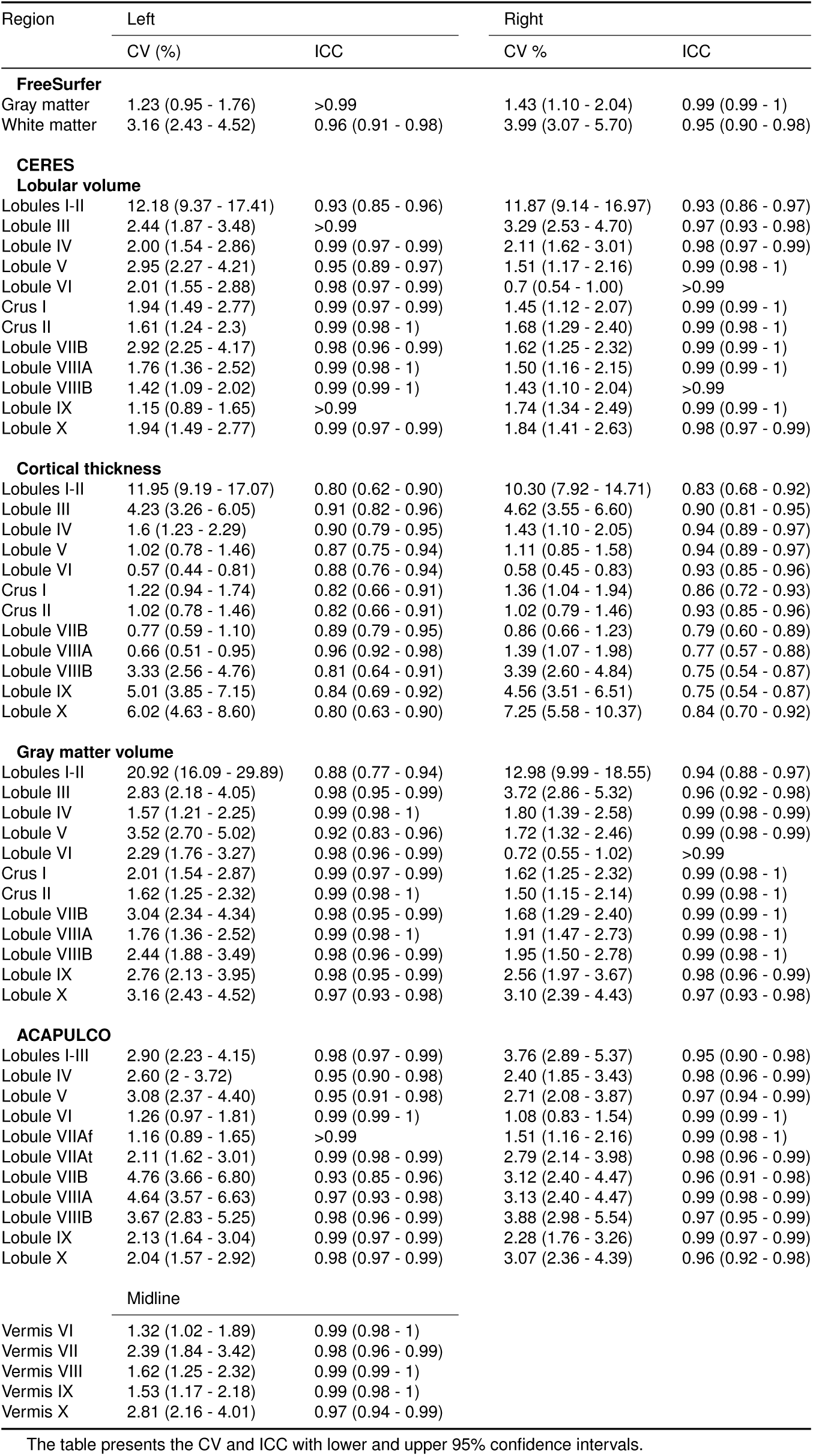
Coefficient of variation (CV) and intraclass correlation coefficient (ICC) for cerebellar regions obtained by FreeSurfer, CERES, and ACAPULCO with data from the Kirby-21 study.

### Visual inspection

All fully automated analyses resulted in anatomically broadly correct segmentations and parcellations except one FreeSurfer analysis (analysis failure) and one ACAPULCO analysis (incorrect placement of the bounding box localizing the cerebellum). In several FreeSurfer analyses, voxels containing dura and surrounding non-brain tissue were mislabeled as cerebellum, in particular, in the midline. In single ACAPULCO analyses, the parcellation algorithm mislabeled voxels located in the neck as cerebellum, even after postprocessing (e.g., the second examination of participant OAS2_0013 in the OASIS-2 study). Visual inspection of all CERES analyses did not reveal remarkable inaccuracies.

### Replicability

Using FreeSurfer with data from the ChroPain2 study, two identical analyses of the same T1-weighted image provided identical results for all participants regarding gray and white matter volumes.

Using CERES with data from the ChroPain2 study, two identical analyses provided identical results for lobular volumes, cortical thickness, and gray matter volumes in most participants (Figure 4D, E, F for right lobules V, VI, VIIIA). For lobular volumes, differences for all regions were smaller than ± 0.1%. For cortical thickness, maximum differences were found in the left lobules I-II (−4.8 – 3.7%). Maximum differences in gray matter volume were also found in the left lobules I-II (−5.8 – 11.8%, data not shown).

Using ACAPULCO with data from the ChroPain2 study, two identical analyses provided different results for all regions. Differences were larger than those found with CERES (Figure 4B). Differences were between -36.6% (right lobule VI-IIB) and 20.6% (right lobule IX). To confirm these results, analysis replicability of ACAPULCO was also assessed with all T1-weighted images of the Kirby-21 study (Figure 4C). For this data set, differences were between -11.5% (vermis X) and 19.4% (left lobule VIIIB, data not shown).

### Repeatability

Comparing the FreeSurfer results of the first and second T1-weighted MRI in the Kirby-21 study, differences in gray matter volumes were below ±5% (Figure 5A). Differences in white matter volumes were higher, between −12.1% and 19.2%. With CERES, differences in lobular volumes, cortical thickness, and gray matter volume were below ±5% in most cases (Figure 5C, D, E). In some cases, differences were considerably higher, in particular for the small lobules I-II. With ACAPULCO, differences in lobular volumes were also below ±5% in most cases (Figure 5B). Maximum differences were between -20% (left lobule VIIB) and 35.1% (left lobule VIIIA, data not shown).

The image intraclass correlation coefficients (I2C2) for repeatability using the CERES parcellations are presented in Table 2. The coefficients of variation and the intraclass correlation coefficients for repeatability are presented in Table 3. Most lower 95% confidence intervals suggest good or even excellent repeatability.

### Long-term reproducibility

Comparing the FreeSurfer results of the first and second T1-weighted MRI in the OASIS-2 study, most differences in gray matter volume were below ±5% (Figure 6A). Maximum differences in gray matter volume were between -12.3% and 9.7%, in white matter volume between -15.6% and 25.5%. With CERES, most differences for lobular volumes, cortical thickness, and gray matter volumes were below ±10%, many even below ±5% (Figure 6C, D, E). Maximum differences for lobular volumes (−60.1%, 167.5%), cortical thickness (−54.4%, 190.8%), and for gray matter volumes (−55.3%, 139.5%) were considerably higher. With ACAPULCO, differences were also below ±10% in the majority of cases, many even below ±5% (Figure 6B). Maximum differences were between -96.4% and 180.9%.

The image intraclass correlation coefficients using the CERES parcellations suggest moderate reproducibility (data not shown). The coefficients of variation and the intraclass correlation coefficients for reproducibility are presented in Table 4.

**Table 4.**
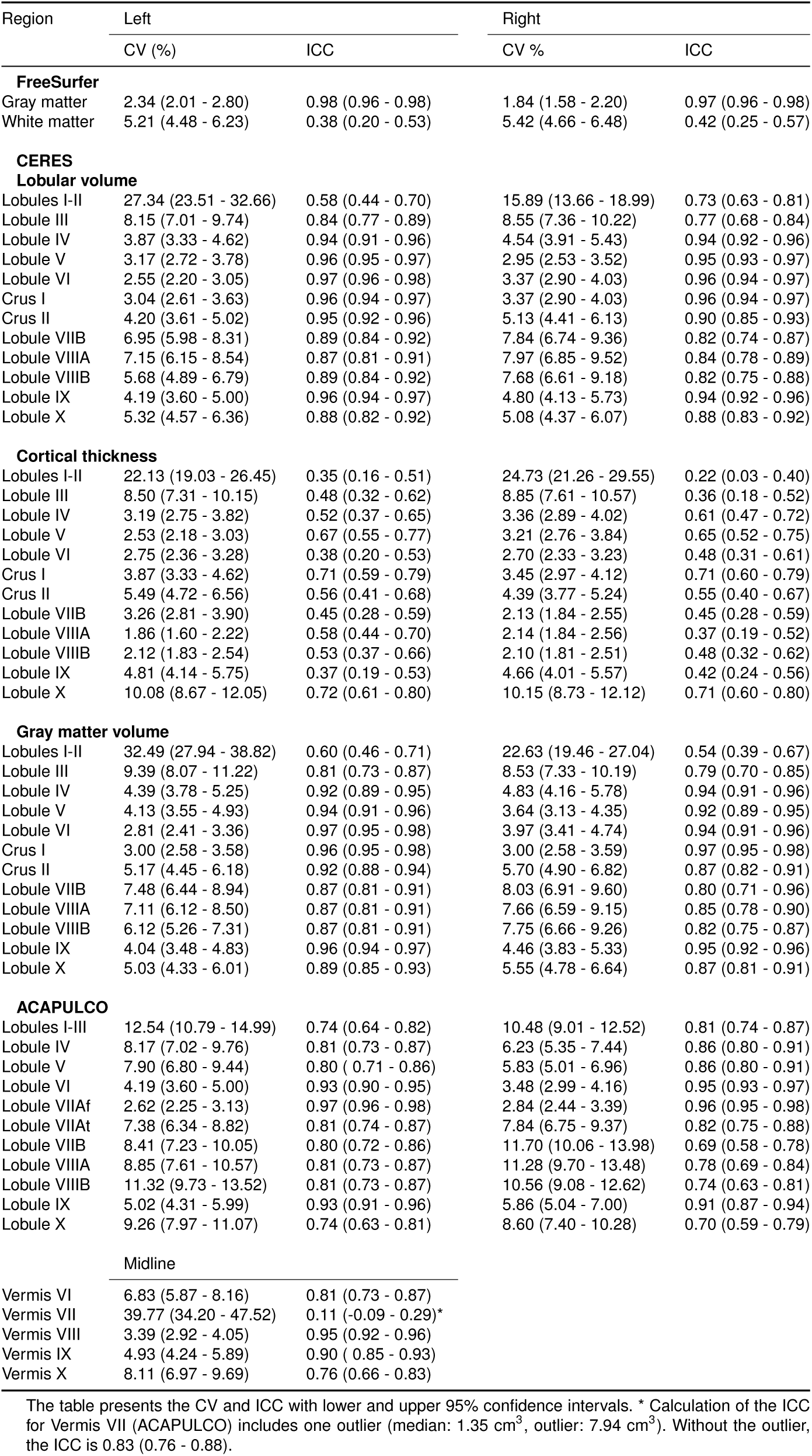
Coefficient of variation (CV) and intraclass correlation coefficient (ICC) for cerebellar regions obtained by FreeSurfer, CERES, and ACAPULCO with data from the OASIS-2 study.

## Discussion

We present a detailed analysis of the reproducibility of fully automated cerebellar morphometry using three different software packages regarding (1) replicability (two analyses of one data set with identical hardware and software), (2) repeatability (analyses of two data sets taken on the same day), and (3) long-term reproducibility (analyses of two data sets taken months or years apart).

Regarding analysis replicability, we found that the results of FreeSurfer segmentations were identical in all analyses. Replicability was high for CERES parcellations and segmentations in most regions (Figure 4D-F), although the Patch-Match algorithm employed by CERES is non-deterministic and involves a random search step that is performed iteratively (20). By contrast, we found substantial differences when performing two identical ACAPULCO analyses of the high-resolution ChroPain2 data sets (Figure 4B). We hypothesized that the submillimeter resolution (0.75 mm isotropic voxel size) of this data set might have caused problems for ACAPULCO’s parcellating network which has been trained with MPRAGE images resampled to 1 mm isotropic resolution (24). Therefore, we assessed ACAPULCO’s analysis replicability with data from the Kirby-21 study (1 × 1 × 1.2 mm^3^ voxel size). Differences between two identical ACAPULCO analyses were lower in the Kirby-21 study compared to the ChroPain 2 study (Figure 4B-C) but still relatively high, with most differences <±5%.

Assessment of repeatability revealed a remarkably similar picture for all software packages (Figure 5). Most differences between the first and the second MRI taken on the same day were <±5%. This result presents an estimation of the reproducibility with which cerebellar subdivisions can be determined with a recent MRI system at 3 Tesla, a widelyused MPRAGE sequence, and a fully automated segmentation and/or parcellation software for individual participants today. For ACAPULCO, intraclass correlation coefficients in our study using the Kirby-21 data set (Table 3) were similar to the ICCs reported in Han et al. (24), although both studies used different algorithms.

For comparison, estimation of cerebral cortical thicknesses using FreeSurfer demonstrated an overall higher reproducibility with differences between scans taken within minutes of ≤±1.9% and between scans taken within weeks of ≤±2.3% (56). Of course, the reported differences between two scans of one person is a complex mixture of several factors, including not only imperfections of the image analysis software used, but also of scanner hardware and MRI sequences, and differences in the positioning of the head. Using a high-resolution sequence (e.g. with a 0.75 mm isotropic voxel size) and/or a higher magnetic field strength (i.e., 7 Tesla) is expected to improve not only assessment of cerebral cortical thicknesses (57) but also of cerebellar volumes and cortical thicknesses due to reduced partial volume effects or increased signal-to-noise-ratios. As the developers of CERES acknowledge, the main limitation of their analysis software is the small library of only five manually labeled cerebellar templates on which CERES relies at present (20). Hopefully, the developers will include additional templates in future versions of their software, likely improving segmentation and parcellation results.

As expected, long-term reproducibility of cerebellar morphometry was lower than repeatability on the same day. Brain volumes and cortical thicknesses change over time, not only due to aging, but also due to factors unrelated to aging, such as diurnal factors (58), hydration (59), or alcohol intake (60). In single cases, both CERES and ACAPULCO analyses resulted in dramatic differences, suggesting mislabeling of large parts of cerebellar regions.

### Recommendations for use of automated cerebellar morphometry

Based on the presented analyses, we recommend the following steps to improve the design, data analysis, and interpretation of future neuroimaging studies:

**1. Quality control through visual inspection of all labeled regions**. Corroborating the results of Kavaklioglu et al. (61), we recognized that FreeSurfer frequently mislabeled voxels representing the dura mater or the dural sinuses as cerebellar gray matter. The number of these voxels is usually small compared to the entire gray matter of the left or right cerebellum. Manual correction of labels and recomputing of cerebellar volumes is possible, but would require substantial expertise and time (62), and is therefore not feasible in large-scale studies. Of note, the locating network used in ACAPULCO failed in one analysis. In this case, the parcellating network mislabeled all voxels and finished without error message. Thus, we strongly recommend the visual inspection of all results of neuroimaging pipelines, including automated cerebellar morphometry. Visual inspection of subcortical FreeSurfer results requires manual loading of .mgz files in FreeSurfer’s Freeview file viewer or in another viewer capable of displaying .mgz files (e.g., FSLeyes). Visual inspection of CERES and ACAPULCO results is less time-consuming because both analysis packages create report pages in pdf or html format for convenient inspection.

**2. Assessment of analysis replicability**. Many MRI analysis packages include stochastic algorithms, such as random seed generation for the initialization of analyses (63). Given the remarkable differences found in identical analyses by ACAPULCO, we recommend reporting the analysis replicability for every neuroimaging pipeline, including cerebellar morphometry.

**3. Assessment of repeatability**. For cross-sectional studies, we recommend reporting the repeatability of the selected neuroimaging pipeline in addition to its analysis replicability. The data set for assessment of repeatability should include two identical MRI scans taken on the same day, ideally directly one after another, but with repositioning in between, to minimize true changes in brain volumes or cortical thicknesses.

**4. Assessment of long-term reproducibility**. For the design of a longitudinal study, we recommend investigating the long-term reproducibility of the selected neuroimaging pipeline in addition to its analysis replicability. The data set for estimation of long-term reproducibility should include two or more scans, taken in time intervals comparable to the planned longitudinal study. The obtained results should guide the decision if the expected changes may be observed with the sample size and the study design under consideration (64).

### Conclusions

Based on its high accuracy (19), its overall high reproducibility shown here, and its ability to differentiate between entire lobular volumes, gray matter lobular volumes, and lobular cortical thicknesses, CERES is a powerful tool to investigate cerebellar morphometry. Cerebellar morphometry is expected to provide important biomarkers for cerebellar aging and disease. Reliable neuroimaging biomarkers depend on reproducible analyses. For every neuroimaging pipeline, not only for cerebellar morphometry, reproducibility should be investigated, reported, and utilized for the interpretation of its results.

## ACKNOWLEDGEMENTS

The authors wish to thank Stefan Harfst and Fynn Schwietzer, Scientific Computing, University of Oldenburg, Germany, for continuous support with the HPC cluster. The ChroPain2 study was supported by the Neuroimaging Unit, University of Oldenburg, funded by grants from the German Research Foundation (DFG; 3T MRI INST 184/152-1 FUGG and MEG INST 184/148-1 FUGG). The high-performance computer cluster CARL, University of Oldenburg, is funded by a grant from the German Research Foundation (DFG; INST 184/157-1 FUGG).

## Footnotes

1 www.drks.de

2 uol.de/en/medicine/biomedicum/neuroimaging-unit

3 www.nitrc.org/projects/multimodal

4 www.oasis-brains.org

5 uol.de/en/school5/sc/high-perfomance-computing/hpc-facilities/carl

6 https://surfer.nmr.mgh.harvard.edu/fswiki/rel7downloads

7 volbrain.upv.es

8 cobralab.ca/atlases/Cerebellum

9 iacl.jhu.edu/index.php?title=Cerebellum_CNN

10 fsl.fmrib.ox.ac.uk/fsl/fslwiki/InitialProcessing

11 www.nitrc.org/projects/robex/

12 www.bic.mni.mcgill.ca/ServicesAtlases/ICBM152NLin2009

13 www.tensorflow.org

14 https://cran.r-project.org/web/packages/psych/index.html

15 https://neuroconductor.org/package/I2C2

16 https://osf.io/n8y5h/

